# Altered DNA methylation in ion transport and immune signalling genes is associated with severity in Pancreatic Ductal Adenocarcinoma

**DOI:** 10.1101/2021.06.14.448193

**Authors:** Ankita Chatterjee, Akash Bararia, Debopriyo Ganguly, Paromita Roy, Sudeep Banerjee, Shibajyoti Ghosh, Sumit Gulati, Supriyo Ghatak, Bitan Kumar Chattopadhay, Arindam Maitra, Priyadarshi Basu, Aniruddha Chatterjee, Nilabja Sikdar

**Affiliations:** Human Genetics Unit, Indian Statistical Institute, Kolkata, India; National Institute of Biomedical Genomics, Kalyani, India; Department of Pathology & Department of Gastrointestinal Surgery, Tata Medical Center, Rajarhat, Kolkata, India; Department of General Surgery, Medical College and Hospital, Kolkata, India; Department of HPB Surgery, Calcutta Medical Research Institute, Kolkata, India; Department of General Surgery, IPGMER & SSKM Hospital, Kolkata, India; Department of Pathology, Dunedin School of Medicine, University of Otago, Dunedin, New Zealand

**Author notes:** **Correspondence:** NilabjaSikdar, Ramalingawami Fellow/Scientist D, Human Genetics Unit, Biological Sciences Division, 203, B. T. Road Kolkata -700108.

**Keywords:** Pancreatic-Ductal Adenocarcinoma_2_, 450K DNA methylation_3_, transcription_4_, ion-transport pathway_5_, interferon signalling_6_. (Min.5-Max. 8)

## Abstract

**Background:** Pancreatic ductal adenocarcinoma (PDAC) is one of the leading cancers worldwide and has a poor survival, with a relative five-year survival rate of only 8.5%. In this study we investigated epigenetic marksassociated with PDAC severity and prognosis, through studying alterations in DNA methylation patterns.

**Methods:** DNA methylome for tumor and adjacent normal tissue samples from PDAC patients (n=7) were generated using Illumina 450K bead chips. Differentially methylated positions (DMPs) were identified with |delta beta| > 0.2 and p-value<0.01 by comparing tumors with the adjacent normal tissues. Validation of differential methylation and associated gene expression at selected genes was carried out in an independent cohort PDAC patient.

**Results:** We identified 76 DMPs in PDAC patients that mapped to 43 genes. Among them, 44.7% (n=34) were hypo-methylated and 55.3% (n=42) were hyper-methylated DMPs in cancer samples. The trends of change in methylation at these 76 DMPs from well to moderate were like that from moderate to poorly differentiated cancer samples. The gradual trend in differential methylation was observed both in our cohort and the TCGA-PAAD cohort, suggesting methylation marks can serve as early indicators of disease pathology. Altered promoter methylation, which may affect gene expression, was observed for transcription regulators (*BHLHE23, GSC2, FOXE1* and *TWIST1*), gated ion channels (*KCNA6*, and *CACNB2*), tumor suppressors (*RASSF1*, *SPRED2*, and *NPY*) and genes functioning in interferon signalling (*SIGIRR*, *MX2*, and *OAS2*). We also have compared the TCGA-PAAD dataset with normal pancreatic tissue data from GTEx V8 dataset leading to a confluent observation.

**Conclusions:** We reported the first study on methylome in PDAC tumors from patients in India. We identified altered DNA methylation associated with increasing severity in PDAC among some genes like SIGIRR, MX2 along with other previously reported loci. We also concluded a confluence in our observation when comparing the TCGA-PAAD dataset with GTEx V8 dataset.

## 1 Introduction

Pancreatic ductal adenocarcinoma (PDAC) is one of the leading aggressive cancers and is the 4th most common cause for cancer deaths worldwide, with an annual mortality of approximately 350,000 deaths/year(Lucas et al., 2016).According to the GLOBACON 2018 PDAC caused 4, 32,000 death with a incidence of 4, 59,000 (Bray et al., 2018).PDAC has a poor prognosis with a 5-year survival rate of less than 8.5%, despite extensive research on the cancer type (Lucas et al., 2016).Intrinsic heterogeneity across PDAC patients may contribute to poor survival (McGranahan and Swanton, 2017). The absence of prognostic markers to predict subclinical presence of the disease or treatment responses imposes to formulate personalized therapeutic strategies.

Epigenetic changes, i.e., aberrant methylation of DNA, affect gene expression and are important drivers of carcinogenesis (Chatterjee et al., 2018). Altered methylation status at the CpG islands, which mostly locate at the gene promoters and limits the access to transcription factors, significantly affect gene expression. Hyper-methylation at CpG islands with downregulated expressions of tumor suppressor genes (TSG) and hypo-methylation with increased expression of oncogenes have been documented for many cancers (Hansen et al., 2011). Epigenomics has become a promising paradigm for PDAC diagnosis and has identified pathways that can be targeted for therapy (Lomberk et al., 2019).

Past studies had shown that both genetic and epigenetic alterations contributed to PDAC initiation and progression (Biankin et al., 2012; Lomberk et al., 2019; Bararia et al., 2020). PDAC diagnosis based on gene-mutation has only brought partial success in early diagnosis and overall patients’ survival improvement (Lomberk et al., 2019). Distinct epigenetic markers have been observed for specific subtypes of pancreatic cancers and for specific stages of PDAC (Biankin et al., 2012; Mishra and Guda, 2017). Also, studies including pancreatic cancer samples from different populations have identified partially similar methylation marks (Mishra and Guda, 2017; Mishra et al., 2019). The inter-population variation in epigenetic signature may be explained by varied ethnicity, demographic, and occupational factors(Ansari et al., 2016; Storz and Crawford, 2020).

Globally few studies have been done on investigating the changing epigenetic landscapes through progressive stages of PDAC(Nones et al., 2014; Mishra and Guda, 2017; Bararia et al., 2020). In this study, we have reported the altered genome-wide DNA methylation profile of PDAC across progressive stages and compared our results with the TCGA cohort.

## 2 Methods

### 2.1 Study participants

The study was approved by the Institutional Review Board (IRB) of Indian Statistical Institute, Kolkata and all involved hospitals. Our study included samples of PDAC patients, collected from Tata Medical Center (TMC), Rajarhat, Kolkata, Calcutta Medical Research Institute (CMRI), Calcutta Medical College and Hospital, Seth Sukhlal Karnani Memorial Hospital (SSKM) & Institute of Post Graduate Medical Education & Research (IPGME&R) Hospital, Kolkata during the time frame of 2014 – 2020. Tumor and adjacent normal tissue samples were collected during surgery and stored in RNA later solution. Histopathological examination and TNM staging were done according to the 8th edition guidelines by American Joint Committee on Cancer (AJCC), by a single clinician. In the discovery cohort, 3 were poorly differentiated adenocarcinoma (PDA), 2 were moderately differentiated adenocarcinoma (MDA), and 2 were well differentiated adenocarcinoma (WDA) samples (Table 1).

**Table 1:** Demographic and clinical characteristics of the discovery cohort.

| Discovery cohort (n=7) |  |  |  |  |  |
| --- | --- | --- | --- | --- | --- |
| Patient ID | Sex | Age (Yrs) | Tobacco intake | TNM Grade | Histopathological Report |
| PDAC-1 | Female | 48 | no | T2N1M0 | Poorly differentiated |
| PDAC-2 | Female | 74 | no | T3N0 | Well differentiated |
| PDAC-3 | Male | 38 | no | T3N1M0 | Poorly differentiated |
| PDAC-4 | Male | 65 | yes | T3N1 | Moderately differentiated |
| PDAC-5 | Male | 64 | no | T3N1 | Moderately differentiated |
| PDAC-6 | Male | 43 | yes | T3N0 | Well differentiated |
| PDAC-7 | Male | 40 | yes | T3N1 | Poorly differentiated |
| Validation cohort (n=20) |  |  |  |  |  |
| PDAC-8 | Male | 45 | yes | T2N1M0 | well-differentiated |
| PDAC-9 | Male | 48 | yes | T2N1M0 | well-differentiated |
| PDAC-10 | Female | 74 | no | T3N0 | well-differentiated |
| PDAC-11 | Male | 60 | yes | T3N1M0 | Poorly-differentiated |
| PDAC-12 | Female | 60 | no | T3N0 | well-differentiated |
| PDAC-13 | Male | 44 | no | T2N1M0 | Poorly-differentiated |
| PDAC-14 | Female | 35 | no | T3N0 | well-differentiated |
| PDAC-15 | Male | 43 | yes | T3N0 | well-differentiated |
| PDAC-16 | Male | 50 | no | T2N0Mx | well-differentiated |
| PDAC-17 | Male | 65 | yes | T2N1M0 | Moderately-differentiated |
| PDAC-18 | Male | 60 | no | T2N1M0 | Moderately-differentiated |
| PDAC-19 | Male | 58 | no | T2N0Mx | well-differentiated |
| PDAC-20 | Male | 40 | yes | T3N1 | Poorly-differentiated |
| PDAC-21 | Female | 60 | no | T3N1Mx | well-differentiated |
| PDAC-22 | Male | 70 | no | T3N1 | well-differentiated |
| PDAC-23 | Male | 53 | no | T3N1M0 | Moderately-differentiated |
| PDAC-24 | Male | 68 | no | T3N2MX | Poorly-differentiated |
| PDAC-25 | Female | 54 | no | T2N1MO | Moderately-differentiated |
| PDAC-26 | Male | 62 | yes | T3N1M0 | well-differentiated |
| PDAC-27 | Male | 53 | yes | T3N1MO | Moderately-differentiated |
| PDAC-28 | Male | 54 | no | T2N1 | Moderately-differentiated |
| PDAC-29 | Female | 53 | no | T2N1M0 | well-differentiated |

Preliminary study findings on methylation data analysis from the discovery cohort (n=7), were validated on an independent cohort of PDAC patients (n=20).

### 2.2 DNA extraction and bisulfite treatment

Tissue DNA isolation was done using the QIAGEN DNA extraction kit (DNeasy Blood and Tissue Kit, QIAGEN Inc., Germany) kit. The concentrations and the purity of the DNA samples was estimated using NanoDrop 2000 (Thermo Fisher Scientific™, USA). Bisulfite conversion of DNA samples (~500ng) was done using EZ DNA methylation Gold Kit (Zymo Research, USA). Whole genome DNA methylation profiling was done on Infinium HumanMethylation450 BeadChip (Illumina) and signal intensity on bead chips were converted to raw data using the iScan platform (Illumina). All the lab experiments were done following the manufacturer’s protocols. Our final raw methylome data was generated on 7 PDAC patients.

### 2.3 Processing of DNA methylation data

For each CpG site in the genome, methylation level was assessed using the established β value. β value was calculated as the ratio of fluorescent signal intensity of the methylated (M) and total of signal intensities from the methylated (M) and unmethylated (U) alleles:

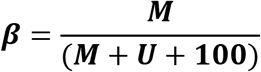

We have used the minfi R package (Aryee et al., 2014) to convert raw array data to β value and for data normalization. To reduce false positive inferences from our data, we have used data quality control as described by Aryee et. al.. Probes were excluded from further analysis if : (1) their detection p-value>0.05, (2) they contain SNPs in their sequences, (3) they are positioned on X and Y chromosomes(Fortin et al., 2014, 2017) (Supplementary figure 1) (Aryee et al., 2014).

### 2.4 Differential methylation analysis

Differentially methylated positions (DMPs) between the tumor and the normal tissue samples were identified using “dmpFinder” function included in the minfi package. Probes were considered as differentially methylated (DM), if the p-value corrected by Benjamini-Hochberg (BH) multiple corrections methods was <0.05 and the absolute value of Δβ (difference between the average β-value among the tumor samples and the average β-value among the normal samples) >0.2. Hyper-methylated positions (CpG sites) were DMPs with Δβ >0.2 and hypo-methylated sites were DMPs with Δβ<−0.2 (Supplementary figure 1).

### 2.5 Comparison with TCGA and GTEx data set

We have compared our results with the methylation data of tumor and solid normal tissues from 9 PDAC patients, including the TCGA database. Change of methylation at DMPs with different pathologic stages were observed using methylation data of 182 additional tumor samples (TNM stage I, n=21; TNM stage II, n=150; TNM stage III, n=5; TNM stage IV, n=4) (Supplementary table 1). Differential methylation analysis between PDAC tumors and normal tissues and between different stages of tumor tissues were done as previously described in the methods 2.4.

Gene expression data (FPKM) was downloaded for the differentially methylated genes for the same set of TCGA samples (n=182). Correlations between methylation statuses at CpG site in a gene with its expression, was estimated. Expression of genes in normal pancreatic tissue samples (n=248), that were differentially methylated, was downloaded from GTEx database V8. Comparison of expression of these genes between pancreatic cancer samples (from TCGA cohort) and normal pancreatic tissues (from GTEx database V8) was performed.

### 2.6 Validation

Differential methylation at the gene promoters is often inversely associated with its expression. Expression of selected genes, where we observed alteration in their promoter methylation, was studied among the tumor and normal tissue samples collected from the validation cohort. An independent validation cohort of 20 PDAC patients was formed for studying promoter associated methylation marks in PDAC. Alteration of promoter methylation was studied using methylation specific PCR (MSP). Bisulfite conversion of DNA samples was done as mentioned earlier in section 2.2. The final DNA concentration was 10ng/μl. The details of the MSP are described in supplementary methods. Differential gene expression was studied in a separate cohort of 20 PDAC patients. Gene expression of the selected genes was estimated using real-time PCR, using iTaq Universal SYBR Green Supermix fluorescent dye (Bio Rad Life Sciences Research). Extended protocol for gene expression study was mentioned in detail in Supplementary methods section.

### 2.7 Functional characterization of the differentially methylated genes

The DMPs were annotated to their nearest genes with reference genome annotation hg19. The identified gene list was designated as genes with differential methylation (gDM) in this article. Enrichment of Gene Ontology (GO) terms on biological processes among the gDMs was done using Metascape. Network construction and protein-protein interactions (PPI), depicting functional relationships among the gDMs, was done using Ingenuity Pathway Analysis (IPA) and Cytoscape, respectively. Significantly enriched GO terms were identified using right-sided hypergeometric tests and multiple-testing correction was done using the Benjamini-Hochberg method. The expression of target genes in pancreatic cancer and their relationship with disease prognosis was observed using Protein atlas database (https://www.proteinatlas.org/). For the genes, where altered promoter methylation was observed, detailed gene functions had been studied through literature search.

### 2.8 Statistical analysis

Differential methylation between cancer and normal samples were studied through linear modelling after adjusting for the effects of age and gender, using the “minfi” R package. Hierarchical clustering method was used depending on the estimated Euclidean distance between the samples to classify the samples. The R functions “hclust” and “heatmap3” were used for heatmap preparation. Relationship of gene expression with disease prognosis were identified using Kaplan-Meir survival analysis, which were done using Protein atlas database (https://www.proteinatlas.org/).

## 3 Results

### 3.1 Characteristics of patients

The demographic and clinical features of the 7 patients recruited in our study are explained in Table 1. Among the 7 patients, 5 were males and 2 were females. Average age of the patients was 53.14 years. Among the males, 3 had a history of tobacco intake, either through smoking or chewing habits. None of the females had any history of tobacco intake. Histopathological analysis showed different grades of tumour among the patients with increasing severity: well differentiated (WDA: n=2), moderately differentiated (MDA: n=2) and poorly differentiated (PDA: n=2) (Table 1).

The validation of methylation was done among 20 PDAC patients. Among them, 20% (n=4) of the patients had poorly differentiated PDAC and 30% (n=6) had moderately differentiated PDAC. 40% (n=8) had a history of tobacco intake (Table 1). The validation of gene expression was done on a separate group of 20 PDAC patients, among whom 21% (n=4) had well differentiated, 47.3% (n=9) had moderately differentiated and remaining 31.5% (n=6) had poorly differentiated PDAC stage.

### 3.2 Differential methylation in PDAC

A list of 2213 DMPs in PDAC, that were previously identified by *Nones et al*. (Nones et al., 2014), was selected and a targeted approach was taken to study the methylation status at established DMPs in PDAC cancers. Given the small sample size in our study, this exploratory analysis was done to identify overall true positive differential methylation signature in PDAC. Hierarchical clustering of the study samples with these 2213 DMPs showed a correct classification of 72% (Supplementary figure 2). So, we concluded that overall the differential methylation signature in PDAC samples were like the previous findings. The misclassification of the remaining 2 samples (28%) might be due to sample specific characteristics.

In our study we identified a total of 76 differentially methylated positions (DMPs, | Δβ| >0.2, p-value < 0.05), mapped to 43 genes in PDAC tumour compared to adjacent normal tissue samples from 7 PDAC patients (Figure 1A, Supplementary table 2 and 3). Among the 76 DMPs, 44.7% (n=34) were hypo-methylated and 55.3% (n=42) were hyper-methylated among cancer samples. Hierarchical clustering of the samples with these 76 DMPs, showed separated clustering of the cancer and the normal samples (Figure 1A). The mean β-values of the hypo-methylated CpG sites in cancer was less than that in the adjacent normal tissue samples (Supplementary figure 3). Similarly, the mean β-values of the hyper-methylated DMPs were higher among the tumour samples than the normal (Supplementary figure 3).

**Figure 1:**
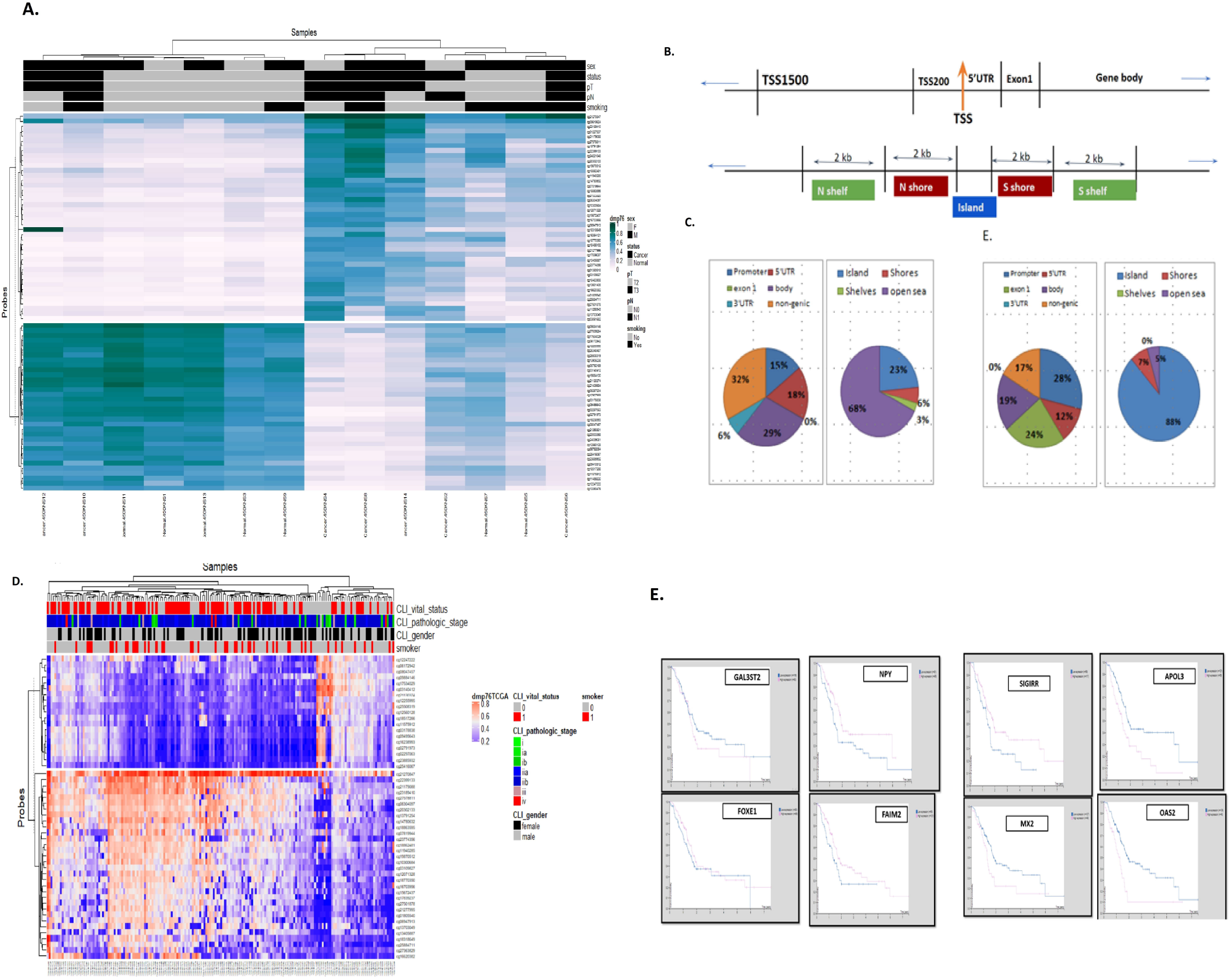
(A) Hierarchical Clustering of the Samples Based on the Top-Ranked DMPs (n=76): Illustrative heat map denoting the Hierarchical Clustering of the PDAC Samples (n=7) based on the Top-Ranked DMPs (n=76). White to yellow to orange denotes increase in beta value (hypermethylation). Clinical information of patients is documented above the cluster along with the CpG site positions documented in the left-hand side. (B) Hierarchical Clustering of the TCGA-pancreatic cancer samples (n=182) on the Top-Ranked DMPs: The columns represented samples and rows represented DMPs (n=76). Clinical phenotypes of patients is documented above the cluster along with the CpG site positions documented in the left-hand side. (C) Genomic annotations of the 450k probes. (D) Genomic annotations of 76 differentially methylated positions (DMPs)-CpG islands (Islands, Shores (±2KB from the boundaries of the islands), Shelves (±2KB from the boundaries of the shores) and Open Sea) or the transcription start site (5’ UTR, Exon 1, Promoter, Body, Non-genic) for Hypomethylated DMPs (n=34) and Hypermethylated DMPs (n=42). (E) Survival plots showing differences in disease prognosis among the TCGA cohort patients with high and low expressions of the gDMs.

Differential methylation signatures in the gene promoters and in the first exon are of noted importance, as previous studies have shown their direct role in regulating gene expression (Brenet et al., 2011; Moarii et al., 2015). Among the hypo-methylated DMPs and the hyper-methylated DMPs, 33% and 64% reside on the gene promoters (1500bp upstream of transcription start site (TSS)), 5’UTR and Exon1, respectively. As was observed in previous studies, hyper-methylation in the cis-gene regulatory segments was more compared to hypo-methylation in the cancer samples (Figure 1B). Aberrant methylation at the CpG islands and surrounding shores (±2kb from CpG islands) were found to be associated with tissue specific and cancer specific methylation signature (Esteller, 2002; Irizarry et al., 2009). Among the hyper-methylated DMPs, 88% resided on the annotated CpG islands and 7% on the CpG shores. Comparatively, hypo-methylated DMPs were less common in the CpG islands (28%) and shores (6%) (Figure 1C). This trend of differential methylation signature was also previously shown in PDAC (Nones et al., 2014).

We then compared the methylome data of PDAC tumour and solid matched tissue normal samples from 9 patients from TCGA database. A total of 7832 DMPs (| Δβ| >0.2, p-value < 0.05) were identified between the tumour and normal samples in the TCGA cohort (Supplementary figure 4A). Among them 54% were hyper-methylated and 46% were hypo-methylated. The 7832 DMPs could correctly classify the cancer and normal samples in separate clusters (Supplementary figure 4B), which indicated the success of our analysis pipeline. Among these 7832 DMPs, 38 DMPs overlap with DMPs, identified from our data (Supplementary figure 4C). The beta values at these 38 DMPs in the TCGA data (for 9 PAAD patients) showed positive correlation with the beta values in our study data (Supplementary figure 4D). The remaining DMPs identified in our study (n=38), which did not overlap with the DMPs identified from TCGA cohort, were novel methylation signatures unique to PDAC in Indian population. Hierarchical clustering of the TCGA pancreatic cancer samples (n=182) with the 76 DMPs showed that the hypo-methylated and hyper-methylated probes identified in our study, showed similar methylation patterns in the TCGA dataset (Figure 1D).

We have further checked the survival of the patients with different expressions of methylated genes in the Pancreatic cancer TCGA cohort using the Protein Atlas data portal. Our hypothesis was that genes with hypo-methylation will have higher expression and *vice versa*. Survival of the patients with high expression for genes, with promoter hypo-methylation, was compared with the patients with low gene expression (Figure 1E). Similarly, survivals of the patients with low expression for genes, with promoter hyper-methylation, were compared with the patients with high gene expression (Figure 1). Our results showed that for four genes with hypo-methylation (*SIGIRR, APOL3, MX2* and *OAS2*), survival of the patients with higher expression were lower than those with lower expression whereas the reverse scenario was observed for the hyper-ethylated genes (*GAL3ST2, NPY, FOXE1* and *FAIM2)*.

### 3.3 Change of methylation status at the DMPs with cancer stage

Previous studies on different cancers have shown, methylation patterns can define cancer stages (Hinoue et al., 2012; Zouridis et al., 2012). We hypothesized that differential methylation signatures are related to increasing severity of PDAC cancers. We observed the degree of differential methylation at 76 DMPs from well differentiated to moderately differentiate to poorly differentiated forms of PDAC samples. A gradual trend of increasing methylation at the hyper methylated sites and decreasing trend of methylation at the hypo methylation sites were observed in our study samples (Figure 2A). The maximum difference (Δβ) was observed between the well differentiated forms and the poorly differentiated cancer forms. These observations may suggest that the DMPs may act as markers of disease progression. For each of the 76 DMPs, average beta value was estimated for each cancer stage in the TCGA cohort and the differences in the mean beta values were plotted (Stage III+IV, II and I) (Figure 2B). The results confirmed a similar trend of methylation changes at the target CpG sites (n=76) with increasing cancer severity, among the samples in TCGA cohort (Figure 2B). We next checked the expressions of genes with sequential methylation changes in the TCGA cohort. For genes which showed sequential hyper-methylation with advanced cancer stage (*SIGIRR, MX2, RASSF1, APOL3, ABHD8),* decreased expression was observed in cancer stage IV compared to stage I (except for *SPRED2*) (Figure 2C). The reverse scenario was observed for some genes (*SLITRK3, FOXE1, TWIST1*) showing sequential hypo-methylation, but not all (Figure 2D). However, when the correlation between methylation and gene expression was observed in the TCGA cohort, we found negative correlations for 60% of the gDMs. (Figure 2E). Thus, we conclude that gradual increase in methylation was observed with increasing cancer severity for hypo-methylated gDMs and *vice versa*.

**Figure 2:**
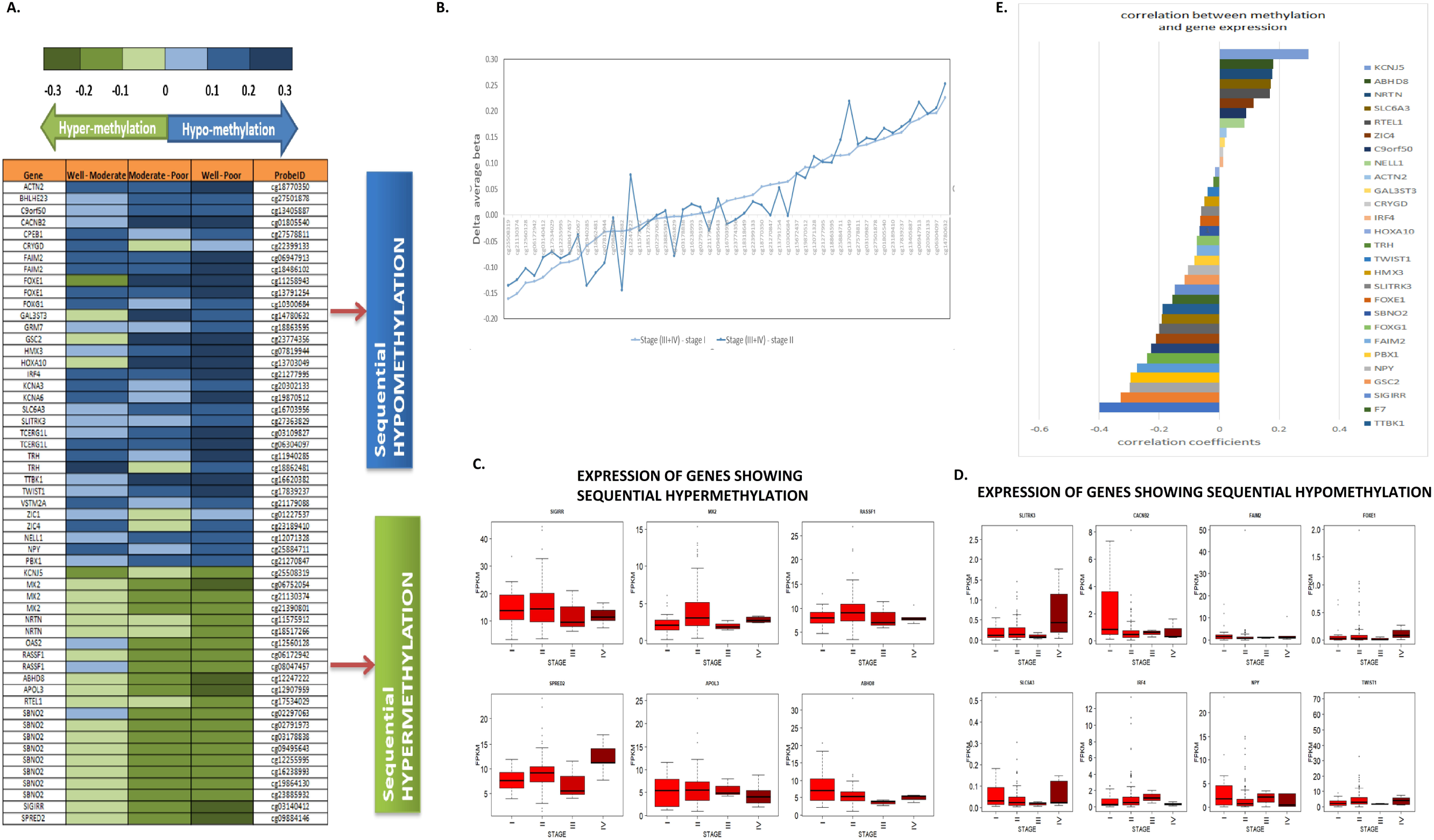
Changes of methylation with increasing severity in the cancer phenotype. (A) Heat map representing differential methylation pattern modulation amongst PDAC associated genes based on varied disease severity. Intensity changes in Blue colour signifies hypomethylation signal dysregulation from Well to moderate to poorly differentiated cancer phenotypes and green colour represents the sequential changes in hypermethylation. Green indicates sequential hypermethylation and blue indicates sequential hypomethylation from well to moderate to poor phenotypes. (B) Changes in methylation at the DMPs from mild to severe stages of cancer in the TCGA cohort. (C) Expression of genes (FPKM) which showed sequential hyper-methylation in our data, was observed across stages of pancreatic cancer in TCGA cohort. (D) Expression of genes (FPKM) which showed sequential hypo-methylation in our data, was observed across stages of pancreatic cancer in TCGA cohort. (E) Correlations between methylation and gene expression in the TCGA cohort.

### 3.4 Relation between promoter methylation at DMPs with gene expression

Among the 76 DMPs, 33% of the hypo-methylated sites and 64% of the hyper-methylated sites were annotated to the promoter (Figure 1A). Change in methylation status at the gene promoter is negatively associated with its expression. Hyper-methylation at the promoter can cause DNA compaction, giving limited access to the binding sites of the transcription factors and *vice versa*. Consequently, promoter hyper-methylation can cause reduced gene expression and *vice versa*. To test this hypothesis in our data, we first studied the correlation between methylation statuses at DMPs annotated to gene promoters with its expression, in the TCGA cohort. Significant negative correlation between gene expression with promoter methylation were observed for *FOXE1* (cg13781254; correlation coefficient=−0.2834, p-value=4.9E-04), *MX2* (cg21130374; correlation coefficient=−0.5, p-value= 5.8E-11) and *CPEB1* (cg27578811; correlation coefficient= −0.54, p-value= 5.8E-11) genes (Figure 2E).

We further extended our search and observed differential methylation in cancer and normal samples, at all the CpG sites annotated to the gene promoters, which contained the identified DMPs. Density plots showed differences in average beta values at each CpG site in the promoters of six genes (*FAIM2, NPY, GSC2, BHLHE23, SLITRK3* and *FOXE1*). Validations of hyper-methylation of *FOXE1* and *NPY* in PDAC samples were done using MSP (Supplementary figure 5). To validate promoter associated methylation, we designed primers for MSP targeting the promoter regions of three selected genes – *FAIM2, NPY, FOXE1.* Data on MSP from the validation cohort (n=16) showed increased % of promoter methylation among the cancer samples compared to the normal samples (Figure 3A). Expression changes of selected gDMs with differential promoter methylation (seven genes selected: *SIGIRR, KCNA6, RASSF1, FAIM2, NPY FOXE1, SLITRK3*) were observed in an independent cohort of 18 paired samples containing tumor and adjacent normal along with 2 unpaired samples containing only tumor specimens. We observed upregulated expressions of hypo-methylated gDMs (*SIGIRR, KCNA6, RASSF1*) in PDAC samples in our validation cohort compared to normal samples (Figure 3B). We further validated our study finding on a larger sample set. We downloaded expressions of gDMs (n=37, for 5 genes expression data was not available) on normal pancreas tissue (n=248) from GTEx database V8. Many of the gDMs were found to be expressed minimally or not at all in the healthy pancreatic tissue samples. Whereas genes that showed prominent expressions were *ABHD8, APOL3, CACNB2, FAIM2, KCNJ5, MX2, NPY, NRTN, PBX1, RASSF1, SBNO2, SIGIRR, SPRED2, RTEL1)* (Figure 3F). We then compared the normal tissue expression data with the expression data of these 37 gDMs in the TCGA cohort. Significant changes in expressions of many genes were observed in pancreatic cancer samples (Figure 3F). Expressions of gDMs (n=37) were significantly higher in the TCGA cohort compared to the GTEx cohort samples.

**Figure 3:**
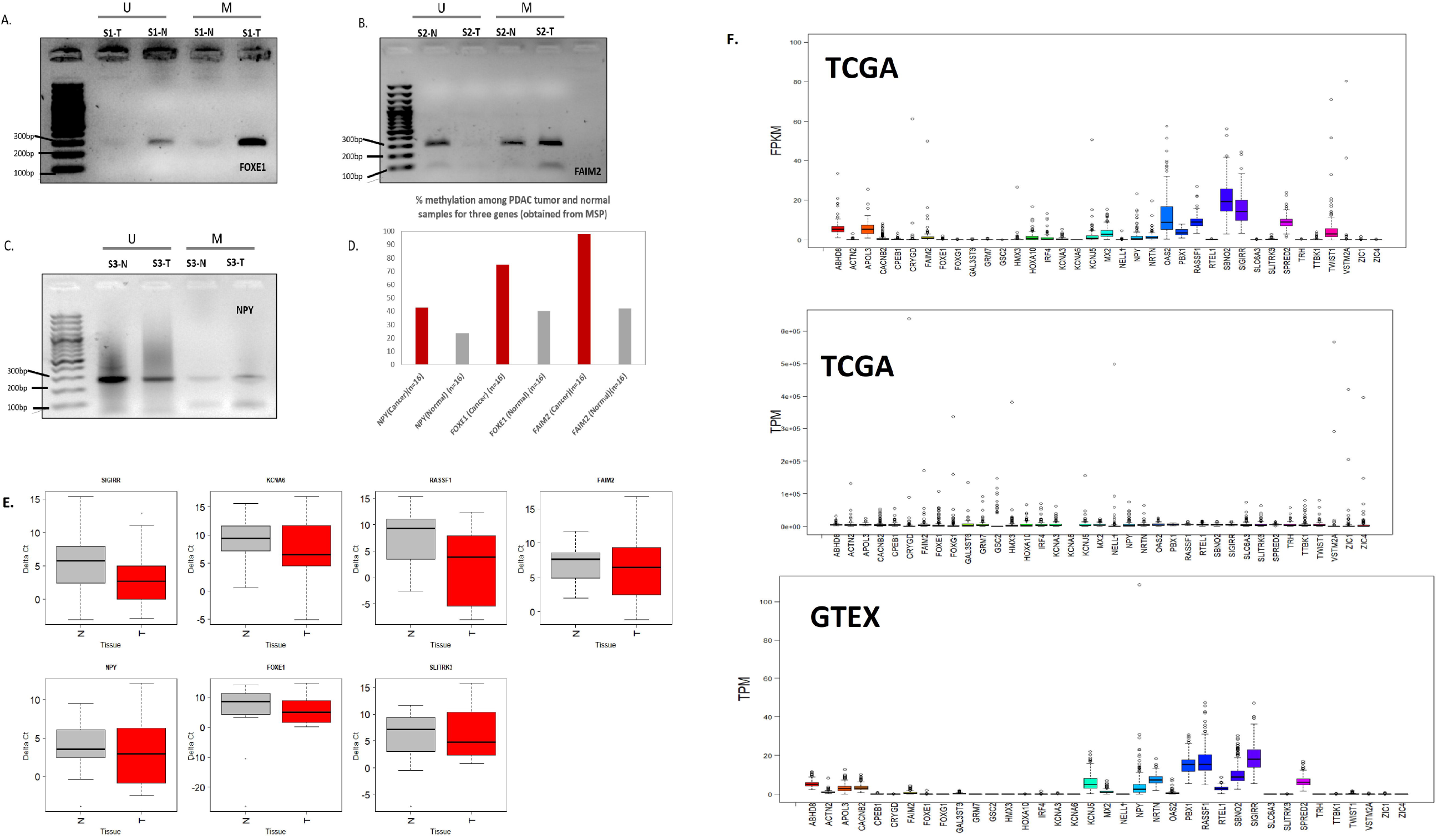
Validation of promoter methylation and its association with gene expression. (A) Representative MSP for *FOXE1.* (B) Representative MSP for *FAIM2.* (C) Representative MSP for *NPY.* (D) % methylation among PDAC tumor and normal samples for three genes. (E) Expressions of seven genes among the cancer and normal samples in the validation cohort. (F) Boxplots showing expressions of gDMs among the pancreatic cancer samples in TCGA cohort (FPKM and TPM, estimated from FPKM) and normal pancreas tissue samples from GTEx database V8.

### 3.5 Molecular significance of differential methylation in PDAC pathogenesis

The 76 DMPs identified in our study were annotated to 43 genes (gDMs) (Supplementary table 2 and 3). Network analysis showed several interactions among the 43gDMs identified in our study (Figure 4, Supplementary table 5). Network analysis with IPA identified three top functional gene groups, which showed differential methylation in PDAC- “Cell Cycle, Gene Expression”, “Organismal Injury and Abnormalities” and “Molecular Transport, Cell-To-Cell Signalling and Interaction” (Figure 4A, 4B, 4C, Supplementary table 5). We have independently constructed the PPI network using STRING database and Cytoscape and identified three main interactive clusters-Cluster 1 (C1) involved in immune regulatory functions, Cluster 2 (C2) involved in ion transport functions and Cluster 3 (C3) included transcription factors and other oncogenic proteins (Figure 4D). Enrichment of biological processes like “regulation of ion transport”, “regulation of response to DNA damage stimulus” and “response to hypoxia”, that were known to be involved in cancer development, were observed among the 43 genes using Metascape (Supplementary table 6). Enrichments of biological processes involved in interactions between cancer and immune system (“Cytokine Signalling in Immune system” and “Interferon alpha/beta signalling”), were also observed in our gDMs. (Supplementary table 6).

**Figure 4:**
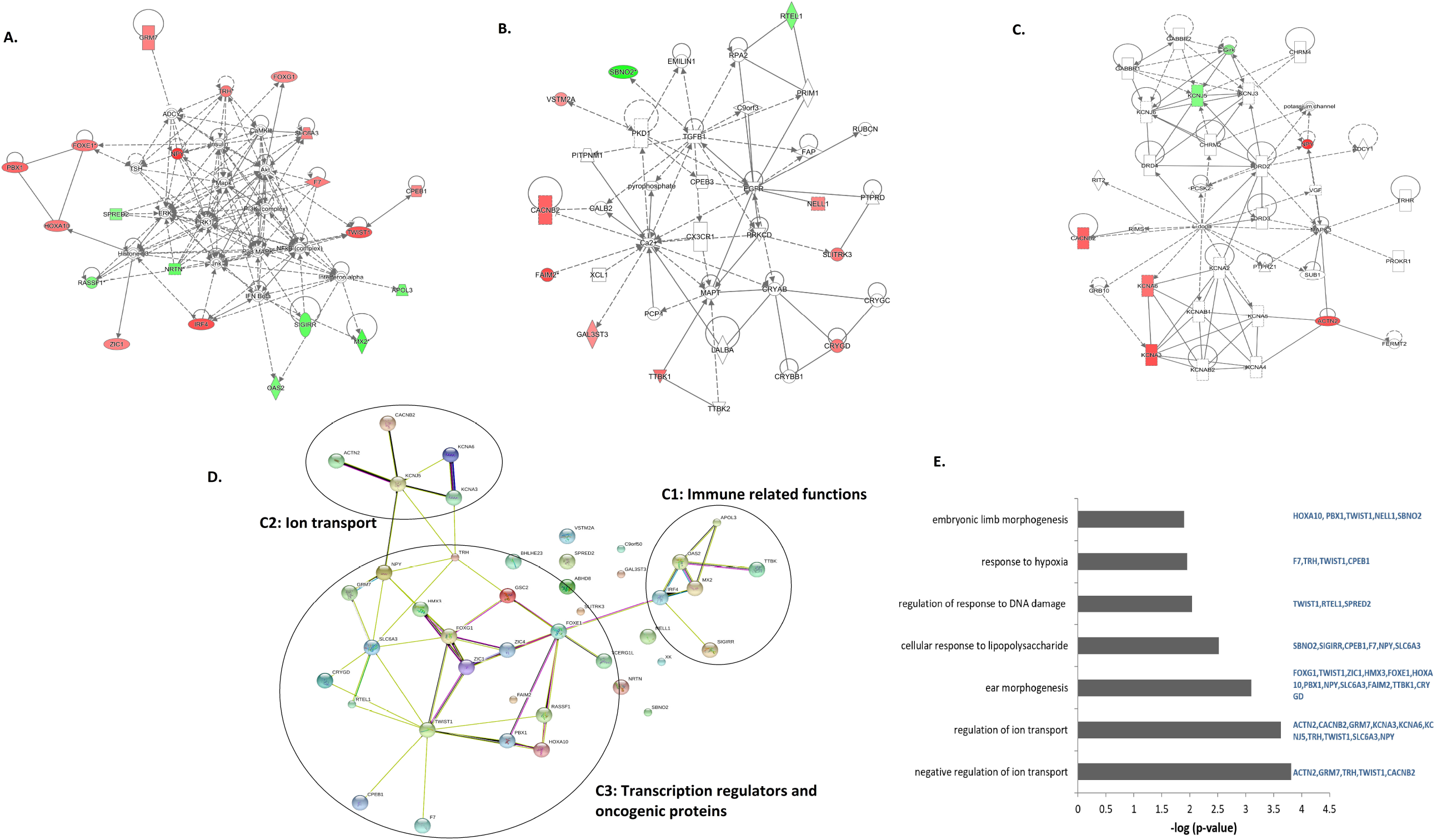
Network analysis done with IPA. Top three networks were shown: (A) Cell Cycle, Gene Expression, Endocrine System Development and Function (B) Developmental Disorder, Ophthalmic Disease, Organismal Injury and Abnormalities and (C) Molecular Transport, Cell-To-Cell Signaling and Interaction. (D) Protein-protein interactions between the gDMs using STRING database. (E) Enrichment of molecular functions among gDMs using Metascape.

## 4 Discussions

Pancreatic cancer, especially ductal adenocarcinoma, is one of the most prevalent cancer types. Late diagnosis, treatment failure, and loco-regional recurrences contribute to its poor prognosis (Ansari et al., 2016). Along with genetic alterations, studies in the past decade showed that epigenetic alterations contribute to PDAC development and pathogenesis (Mazor et al., 2016, Mishra and Guda, 2017; Gregório et al., 2020). Studies also showed that precancerous lesions of PDAC harbour DNA methylation alterations in the known epigenetic driver genes (Bararia et al., 2020). Unusual gene expression is not always due to change in coding sequence but is also related to irregular CpG island methylation profile developed in the DNA’s regulatory region (Hervouet et al., 2018). In regards to varied methylation profile observed amongst various cancer types, a specific abnormal methylation profile can serve as a biomarker for that particular cancer type (Thompson et al., 2015). Since methylation marks can be reversed to an un-methylated state, epigenetic marks can act as a lucrative therapeutic target (Baylin and Jones, 2011). Decrease of overall methylated-Cytosines and increased methylation in normally unmethylated CpG sites, especially in the promoters of tumour suppressor genes, DNA repair genes are observed in tumours compared to normal tissues, which aid in cancer progression and metastasis (Feinberg and Vogelstein, 1983; Gama-sosa et al., 1983; Herman et al., 1994). Recent findings showed abnormal DNA methylation can mark the spectrum of cancer progression, thus serving as biomarkers for diagnosis and prognosis (Dumitrescu, 2018; Vedeld et al., 2018; Peng et al., 2020). In this study we identified DNA methylation alterations associated with increased severity of PDAC samples, which may play a key role in disease progression.

Similar to previous findings (Nones et al., 2014; Mishra and Guda, 2017; Gregório et al., 2020), we have also observed that the majority of the DMPs were hyper-methylated. Hyper-methylation at CpG island in promoters is associated with gene silencing during tumorigenesis, specifically by reduced expression of tumor suppressor genes. In our exploratory analysis, we had observed hyper-methylation in the promoters of transcription regulator genes (BHLHE23, GSC2, FOXE1 and TWIST1). Previous studies have reported hyper-methylation of these transcription regulators in multiple cancers including pancreatic cancer (Vincent et al., 2011; Kinugawa et al., 2017; Natale et al., 2019) and reduced expression of these genes were associated with worse prognosis of PAAD cancer in the TCGA cohort. FOXE1 is one of the most frequently hyper-methylated tumor suppressor genes (TSGs) in pancreatic cancer (Kinugawa et al., 2017; Natale et al., 2019). Hyper-methylation of another previously reported TSG, TCERG1L was observed in PDAC compared to normal tissues (Vincent et al., 2011; Mishra and Guda, 2017). FOXE1, TWIST1 and other hyper-methylated genes involved in cell cycle regulation were observed in PDAC cancers, thus suggesting that their reduced expression may encourage uncontrolled cell proliferation in PDAC.

Promoter hyper-methylation and reduced expression was also observed for SLITRK3 in PDAC. According to Nones et. al., SLIT-ROBO signalling pathways inactivation by epigenetic modulation plays a significant role in activation of HGF/MET as well as in WNT signalling, referring key role in carcinogenesis (Nones et al., 2014). Biankin et. al. reported that out of 58 whole exome tumor data available in ICGC 48% showed hyper-methylation of genes namely, SLIT2, SLIT3, along with ROBO1 and ROBO3. These four genes down-expression associates with SLIT-ROBO pathway inactivation in PDAC (Biankin et al., 2012; Nones et al., 2014).

Another hyper-methylated gene in PDAC was a neurotransmitter Neuropeptide Y (NPY), which can negatively regulate cell motility in cancer by activating the MAPK pathway and regulating intracellular ion channels. NPY is frequently hyper-methylated in certain carcinomas and gene promoter hyper-methylation is correlated with inactivation of gene expression (Ruscica et al., 2007; Li et al., 2015; Bai et al., 2020).

Ion transporters maintain membrane potential and help the cell in vital functions including cell division and obtaining nutrition. Calcium ions also played significant roles in intracellular cell signalling. Deregulations of ion transporters (Calcium, Potassium and Sodium) were observed in many cancers including pancreatic cancers and often represented as biomarkers (Anderson et al., 2019; Patel et al., 2019). Our study first suggests that gradual methylation alterations in several ion transporter genes were observed from early to late stages of PDAC. Ion transporters can help in angiogenesis and cancer metastasis (Munaron, 2015). We observed hyper-methylation of potassium (KCNA3, KCNA6) and calcium (CACNB2) ion channels in PDAC. Studies on animal models showed that epigenetic modification of Kcna3 gene limits T-cell activation (Kang et al., 2016). The voltage gated ion channel Kv1.3 encoded by KCNA3 is activated by changing membrane polarization. The calcium and potassium ion channels thus can inhibit T-cell activation and proliferation upon antigen recognition, creating an immunosuppressive or immune evasive tumor microenvironment in PDAC, which is again associated with poor prognosis and recurrence. Hyper-methylation of potassium channels may thus alter the immune context in PDAC and limit antitumor immunity (Patel et al., 2019).A recent study documented epigenetic dysregulation (hyper-methylation) of Calcium ion transporters in PDAC (Gregório et al., 2020). This study had further shown that epigenetic modification in Calcium signalling occurred at early stages of PDAC.

Promoter hypo-methylation is associated with increased gene expression. In our study promoter hypo-methylation was observed in cell cycle regulatory genes like RASSF1, MX2, SIGIRR, SPRED2. RASSF1 is a potent TSG and was found to be upregulated in Pancreatic cancer (Amato et al., 2016; Malpeli et al., 2019). SPRED2 is another TSG, which negatively regulates Ras-MAPK signalling pathways and its upregulation was related to worse prognosis in pancreatic cancers amples of the TCGA cohort (Jiang et al., 2016).

SIGIRR, MX2, and OAS2 are also involved in Interferon signalling. A recent study showed that overexpression of MX2 reduced cell proliferation, migration, and invasion via ERK/P38/NF-κB pathway in glioblastoma cells (Wang et al., 2019). Upregulated expression of OAS2 was previously reported in pancreatic cancer (Kim et al., 2018; Lv et al., 2019). Interferon induced cell killing is a potent anti-tumor immune response in cancer. A recent study had shown that induction of Type Interferon signalling made pancreatic cancer vulnerable to innate immune systems and showed better response with immune checkpoint therapy (Zhang et al., 2019). Interferon signalling inhibits cell proliferation and cell migration in pancreatic cancer (Fujisawa et al., 2020). Promoter hypomethylation driven upregulated expression of these genes, may thus signify the active antitumor immune response in PDAC. Finally, this has to be hypothesized that Epigenetic and Genetic signature of PDAC partially varies across the globe based on expression frequency and alternated driver behaviors. According to our previous study, Saha et. al., explained KRAS hotspot mutation was observed in low frequency in Indian PDAC population whereas other countries like USA/Canada/European countries and Australia based studies documented KRAS as a high frequency gene around 90% in the same disease. This partially variable profile may be contributed by the patient pool lead by combined factors such as ethnicity, environmental, lifestyle and occupational risk factors (Saha et al., 2020). Similar observations have also been observed in terms of 450K methylome data from several previous studies. In methylome’s case alternate DMP’s along with some common DMP’s have been observed across the globe contributing to variation in the Methylome profile. Various studies conducted on patient samples and across various datasets showed quite partially contrasting signature profiles which might be caused due to these above reasons. In our study, the comparative analysis of data obtained from 450K and TCGA- PAAD dataset we have shortlisted 7 genes for their expression validation in a specific sample set along with their MSP profiling. The 7 gene set consisted of 3 novel genes with respect to our population namely FAIM2, SLITRK3, NPY and FOXE1, was seen to be previously reported. The other 3 genes RASSF1A, SIGIRR, KCNA6 were chosen based on their involvement in 3 different crucial pathways. The expression data obtained showed consistent with that the 3 novel genes were truly found novel in our patient sample set also keeping in concurrence with our observations from 450K methylome data. The RASSF1A, SIGIRR, KCNA6 genes were also found to have significant expression which may conclude that their role in the pathways leading to PDAC are true to our sample set. The novelty of the work could also unrevealed Indian patient population specific differential methylated genes as well as explored unknown epigenetically regulated pathways. Hence these pathways can play a crucial role as therapeutic targets. According to Wang et. al., methylation modulated genes can influence cancers and this varies from country to country (Mishra et al., 2019). In our study also we have observed similar variation in 450K methylome data which can be hypothesized that the different architecture in methylation landscape contributed by factors such as ethnicity, occupational, environmental, lifestyle and demographic (Delpu et al., 2011).

Pancreatic cancer is an overly aggressive in nature with significant difficulty in diagnosis it in early stages due to substantial lack of biomarkers along with the reason of late manifestation of symptoms in PAC patients (Tan et al., 2009). In due respect with PAC aggressive nature and its associated high mortality rate, the urgency of novel therapeutic strategies to combat is of priority. Recent advances in methodological technologies in gene methylation analysis have allowed our view to expand from the single gene level. This remains a valid specific candidate gene approach, to the genome-wide scale, along with which it possesses power in its unbiased approach. Several techniques currently used for methylation specific analysis includes procedures like methylation-specific PCR, array methodology, NGS after bisulfite treatment, and 450K methylation array platform (Omura and Goggins, 2009). Synchronous aberrations between genetic and epigenetic scenarios leads to neoplastic transformation along with cancer phenotype determine signatures and its associated clinical symptoms. To control the devastating effect of PDAC on patients lead by poor prognosis, it is very urgent to understand both the genetic as well as epigenetic events occurring in the disease progression from pre-cancerous lesions to PDAC. Despite numerous clinical trials utilizing known targeted agents, the overall advancement made in pancreatic cancer treatment has been relatively modest in comparison to the advancement made in the treatment of other types of tumors. Initially PAC associated pathogenesis was entirely credited to the genetic mutations but in regard to present technological advances in research, the role of epigenetic aberrations in pathways leading to PAC pathogenesis has also been duly noted and recognized. This observation that targeted the importance of alteration in epigenetic pathways leading to aberrant cancer phenotypes showcased the importance of developing epigenetic inhibitors to down regulate this epigenetic lead phenotype changes. Therefore, the exploration of novel agents targeted at certain epigenetically enriched signaling pathways is one of the most important undertakings to improve the outcome of patients with lethal pancreatic cancer. In addition to a need for disease modelling as well as understanding epigenetic pathways, we must improve the pharmacology of the drugs themselves. Although research has shown time and again that epigenetic drug are effective in eliciting a desire cellular responses much remain unknown about the mechanism of action and cellular pathways involved in driving that responses. Our understanding of epigenetically modulated pathways along with its downstream effects is also limited as of now. Since epigenetic alterations function as a means for cellular response to environmental stimuli, they are by nature very dynamic pathways. Furthermore, the underlying mechanism driving the aberrant activation of certain epigenetic regulators is unknown is many cases. Activation or inhibition of these epigenetic pathways could be a result of genetic mutations in the regulators themselves or activation of upstream signaling pathways, increasing the complexity of using targeted inhibitors for therapy. Currently targeted therapeutics is based on genetic understanding, but epigenetic based targeted therapy can be a boon in curing patients since epigenetic events are reversible in comparison to genetic events mostly. DNA methyltransferases, a class of epigenetic drugs, have been showing promising results in preclinical studies along with histone deacetylase inhibitors and Histone acetyltransferase. Many potent inhibitors of DNMTs are readily available and are being widely used in cancer research today. Combination of epigenetic drugs along with existing targeted therapies can increase the overall survival rates in PDAC patients which highlights the importance of our novel 450K methylome study in PDAC patients across our nation (India) (Dawson and Kouzarides, 2012; Neureiter et al., 2014; Lomberk et al., 2016; Matsuoka and Yashiro, 2016; Paradise et al., 2018; Cheng et al., 2019).

## 5 Conclusion

DNA methylome profiling of PDAC tumors (n=7), identified aberrant methylation in 76 DMPs from 43 genes (gDMs). Difference in promoter methylation was observed in PDAC tumors compared to adjacent controls in four genes namely FAIM2, FOXE1, NPY, and SLITRK3. The novelty of the studyshowed that the trend of methylation at the 76 DMPs remained similar across well to moderate to poor PDAC stages, suggesting the association of the methylation marks with increased disease severity. In this study first reported that among the 43 gDMs, TSGs and cell cycle regulatory genes (RASSF1, NPY, FOXE1, and SPRED2), genes functioning as ion transporters (KCNA3, KCNA6, and CACNB2) and genes involved in antitumor immune (SIGIRR, MX2, APOL3,) response from Indian PDAC patients. Comparative analysis between TCGA-PAAD dataset and GTEx dataset V8 showed trends which were in close pattern with the data we obtained after validation of the 450K methylome data.

## 6 Limitations of the study

We declare two major limitations of this study. Sample size of our discovery cohort was low. Although we have validated our finding using TCGA-cohort, GTEx dataset V8, literature searches, previous findings and through a small validation cohort, our findings need to be validated on a larger cohort. Secondly, due to insufficient sample availability, we could not validate the methylation marks and gene expression on same samples.

## 8 Data Availability Statement

The original raw and cured datasets used and/or analyzed during the current study are available from the corresponding author on reasonable request.

## 9 Author Contributions

A.M., Core Technologies Research Initiative (Co-TeRi), National Institute of Biomedical Genomics (NIBMG) carried out the Illumina 450K bead chip experiment. A.C. A.B. and D.G. carried out the gene expression profiling, methylation Specific PCR validation experiments. The data analysis was done by A.C. A.B, and A.C. Clinically confirmed surgical/resected samples along with patient clinicopathological and demographic data and histology reports for the study from different hospitals were provided by B.K.C., S.B., S.G., S.G., S.G., and P.R. gave input in histopathological findings for clinical samples. P.B. helped in resources and for laboratory experiments. A.C., A.B, A.C. and N.S. prepared the manuscript and did the interpretation of the findings. N.S. conceived, designed, edited, prepared and corresponds the manuscript.

## Acknowledgments

Authors also acknowledged the National Institute of Biomedical Genomics (NIBMG) for instrumental support for performing experiments on an outsourcing basis under the Core Technologies Research Initiative (Co-TeRi) Initiative. Authors are also thankful to Dr. Dheeraj Anchlia, Dr. Sahid Khondaker, Dr. Abhisekh Mohata, Dr. Jitesh Midya, Dr. Vinu Shankar, Dr. Shuchismita Chakraborty, Dr. Debtanu Halder and Dr. Nilanjan Ghosh for collecting samples from respective hospitals. Authors specially thank Mr. Gourab Saha, Miss. Suchandra Pal and Mr. Sanjoy Kr. Dey for collecting samples and RNA isolation. Authors are also thankful to Dr. Sudipto Saha, from Bose Institute (BI), Kolkata for support in IPA pathway analysis. The authors thank Dr. Meenakshi Munshi, from Department of Biotechnology, GOI for supporting and processing the fund. Dr. Aniruddha Chaterjee would like to thank the Rutherford Discovery Fellowship (Royal Society of NZ) for supporting his current position. Authors are also thankful to Prof. Bidyut Roy for strategic input and proofreading of the manuscript. N.S. also thanks to Master. Saraswan Sikdar.

## References

Amato, E., Barbi, S., Fassan, M., Luchini, C., Vicentini, C., Brunelli, M., et al. (2016). RASSF1 tumor suppressor gene in pancreatic ductal adenocarcinoma: Correlation of expression, chromosomal status and epigenetic changes. BMC Cancer 16. doi:10.1186/s12885-016-2048-0.

Anderson, K. J., Cormier, R. T., and Scott, P. M. (2019). Role of ion channels in gastrointestinal cancer. World J. Gastroenterol. 25, 5732–5772. doi:10.3748/wjg.v25.i38.5732.

Ansari, D., Tingstedt, B., Andersson, B., Holmquist, F., Sturesson, C., Williamsson, C., et al. (2016). Pancreatic cancer: Yesterday, today and tomorrow. Futur. Oncol. 12, 1929–1946. doi:10.2217/fon-2016-0010.

Aryee, M. J., Jaffe, A. E., Corrada-Bravo, H., Ladd-Acosta, C., Feinberg, A. P., Hansen, K. D., et al. (2014). Minfi: A flexible and comprehensive Bioconductor package for the analysis of Infinium DNA methylation microarrays. Bioinformatics 30, 1363–1369. doi:10.1093/bioinformatics/btu049.

Bai, Y., Wei, C., Zhong, Y., Zhang, Y., Long, J., Huang, S., et al. (2020). Development and validation of a prognostic nomogram for gastric cancer based on DNA methylation-driven differentially expressed genes. Int. J. Biol. Sci. 16, 1153–1165. doi:10.7150/ijbs.41587.

Bararia, A., Dey, S., Gulati, S., Ghatak, S., Ghosh, S., Banerjee, S., et al. (2020). Differential methylation landscape of pancreatic ductal adenocarcinoma and its precancerous lesions. Hepatobiliary Pancreat. Dis. Int. 19, 205–217. doi:10.1016/j.hbpd.2020.03.010.

Baylin, S. B., and Jones, P. A. (2011). A decade of exploring the cancer epigenome-biological and translational implications. Nat. Rev. Cancer 11, 726–734. doi:10.1038/nrc3130.

Biankin, A. V., Waddell, N., Kassahn, K. S., Gingras, M. C., Muthuswamy, L. B., Johns, A. L., et al. (2012). Pancreatic cancer genomes reveal aberrations in axon guidance pathway genes. Nature 491, 399–405. doi:10.1038/nature11547.

Bray, F., Ferlay, J., Soerjomataram, I., Siegel, R. L., Torre, L. A., and Jemal, A. (2018). Global cancer statistics 2018: GLOBOCAN estimates of incidence and mortality worldwide for 36 cancers in 185 countries. CA. Cancer J. Clin. 68, 394–424. doi:10.3322/caac.21492.

Brenet, F., Moh, M., Funk, P., Feierstein, E., Viale, A. J., Socci, N. D., et al. (2011). DNA methylation of the first exon is tightly linked to transcriptional silencing. PLoS One 6. doi:10.1371/journal.pone.0014524.

Chatterjee, A., Rodger, E. J., and Eccles, M. R. (2018). Epigenetic drivers of tumourigenesis and cancer metastasis. Semin. Cancer Biol. 51, 149–159. doi:10.1016/j.semcancer.2017.08.004.

Cheng, Y., He, C., Wang, M., Ma, X., Mo, F., Yang, S., et al. (2019). Targeting epigenetic regulators for cancer therapy: Mechanisms and advances in clinical trials. Signal Transduct. Target. Ther. 4. doi:10.1038/s41392-019-0095-0.

Dawson, M. A., and Kouzarides, T. (2012). Cancer epigenetics: From mechanism to therapy. Cell 150, 12–27. doi:10.1016/j.cell.2012.06.013.

Delpu, Y., Hanoun, N., Lulka, H., Sicard, F., Selves, J., Buscail, L., et al. (2011). Genetic and Epigenetic Alterations in Pancreatic Carcinogenesis. Curr. Genomics 12, 15–24. doi:10.2174/138920211794520132.

Dumitrescu, R. G. (2018). “Early epigenetic markers for precision medicine,” in Methods in Molecular Biology (Humana Press Inc.), 3–17. doi:10.1007/978-1-4939-8751-1_1.

Esteller, M. (2002). CpG island hypermethylation and tumor suppressor genes: A booming present, a brighter future. Oncogene 21, 5427–5440. doi:10.1038/sj.onc.1205600.

Feinberg, A. P., and Vogelstein, B. (1983). Hypomethylation distinguishes genes of some human cancers from their normal counterparts. Nature 301, 89–92. doi:10.1038/301089a0.

Fortin, J. P., Labbe, A., Lemire, M., Zanke, B. W., Hudson, T. J., Fertig, E. J., et al. (2014). Functional normalization of 450k methylation array data improves replication in large cancer studies. Genome Biol. 15. doi:10.1186/s13059-014-0503-2.

Fortin, J. P., Triche, T. J., and Hansen, K. D. (2017). Preprocessing, normalization and integration of the Illumina HumanMethylationEPIC array with minfi. Bioinformatics 33, 558–560. doi:10.1093/bioinformatics/btw691.

Fujisawa, M., Kanda, T., Shibata, T., Sasaki, R., Masuzaki, R., Matsumoto, N., et al. (2020). Involvement of the interferon signaling pathways in pancreatic cancer cells. Anticancer Res. 40, 4445–4455. doi:10.21873/anticanres.14449.

Gama-sosa, M. A., Slagel, V. A., Trewyn, R. W., Oxenhandler, R., Kuo, K. C., Gehrke, C. W., et al. (1983). The 5-methylcytosine content of DNA from human tumors. Nucleic Acids Res. 11, 6883–6894. doi:10.1093/nar/11.19.6883.

Gregório, C., Soares-Lima, S. C., Alemar, B., Recamonde-Mendoza, M., Camuzi, D., de Souza-Santos, P. T., et al. (2020). Calcium signaling alterations caused by epigenetic mechanisms in pancreatic cancer: From early markers to prognostic impact. Cancers (Basel). 12, 1–22. doi:10.3390/cancers12071735.

Hansen, K. D., Timp, W., Bravo, H. C., Sabunciyan, S., Langmead, B., McDonald, O. G., et al. (2011). Increased methylation variation in epigenetic domains across cancer types. Nat. Genet. 43, 768–775. doi:10.1038/ng.865.

Herman, J. G., Latif, F., Weng, Y., Lerman, M. I., Zbar, B., Liu, S., et al. (1994). Silencing of the VHL tumor-suppressor gene by DNA methylation in renal carcinoma. Proc. Natl. Acad. Sci. U. S. A. 91, 9700–9704. doi:10.1073/pnas.91.21.9700.

Hervouet, E., Peixoto, P., Delage-Mourroux, R., Boyer-Guittaut, M., and Cartron, P. F. (2018). Specific or not specific recruitment of DNMTs for DNA methylation, an epigenetic dilemma. Clin. Epigenetics 10. doi:10.1186/s13148-018-0450-y.

Hinoue, T., Weisenberger, D. J., Lange, C. P. E., Shen, H., Byun, H. M., Van Den Berg, D., et al. (2012). Genome-scale analysis of aberrant DNA methylation in colorectal cancer. Genome Res. 22, 271–282. doi:10.1101/gr.117523.110.

Irizarry, R. A., Ladd-Acosta, C., Wen, B., Wu, Z., Montano, C., Onyango, P., et al. (2009). The human colon cancer methylome shows similar hypo- and hypermethylation at conserved tissue-specific CpG island shores. Nat. Genet. 41, 178–186. doi:10.1038/ng.298.

Jiang, K., Liu, M., Lin, G., Mao, B., Cheng, W., Liu, H., et al. (2016). Tumor suppressor Spred2 interaction with LC3 promotes autophagosome maturation and induces autophagy-dependent cell death. Oncotarget 7, 25652–25667. doi:10.18632/oncotarget.8357.

Kang, J. A., Park, S. H., Jeong, S. P., Han, M. H., Lee, C. R., Lee, K. M., et al. (2016). Epigenetic regulation of Kcna3-encoding Kv1.3 potassium channel by cereblon contributes to regulation of CD4+ T-cell activation. Proc. Natl. Acad. Sci. U. S. A. 113, 8771–8776. doi:10.1073/pnas.1502166113.

Kim, J. C., Ha, Y. J., Tak, K. H., Roh, S. A., Kwon, Y. H., Kim, C. W., et al. (2018). Opposite functions of GSN and OAS2 on colorectal cancer metastasis, mediating perineural and lymphovascular invasion, respectively. PLoS One 13. doi:10.1371/journal.pone.0202856.

Kinugawa, Y., Uehara, T., Sano, K., Matsuda, K., Maruyama, Y., Kobayashi, Y., et al. (2017). Methylation of tumor suppressor genes in autoimmune pancreatitis. Pancreas 46, 614–618. doi:10.1097/MPA.0000000000000804.

Kuroki, T., Tajima, Y., and Kanematsu, T. (2004). Role of hypermethylation on carcinogenesis in the pancreas. Surg. Today 34, 981–986. doi:10.1007/s00595-004-2858-6.

Li, J., Tian, Y., and Wu, A. (2015). Neuropeptide Y receptors: a promising target for cancer imaging and therapy. Regen. Biomater. 2, 215–219. doi:10.1093/rb/rbv013.

Lomberk, G. A., Iovanna, J., and Urrutia, R. (2016). The promise of epigenomic therapeutics in pancreatic cancer. Epigenomics 8, 831–842. doi:10.2217/epi-2015-0016.

Lomberk, G., Dusetti, N., Iovanna, J., and Urrutia, R. (2019). Emerging epigenomic landscapes of pancreatic cancer in the era of precision medicine. Nat. Commun. 10. doi:10.1038/s41467-019-11812-7.

Lucas, A. L., Malvezzi, M., Carioli, G., Negri, E., La Vecchia, C., Boffetta, P., et al. (2016). Global Trends in Pancreatic Cancer Mortality From 1980 Through 2013 and Predictions for 2017. Clin. Gastroenterol. Hepatol. 14, 1452–1462.e4. doi:10.1016/j.cgh.2016.05.034.

Lv, K., Yang, J., Sun, J., and Guan, J. (2019). Identification of key candidate genes for pancreatic cancer by bioinformatics analysis. Exp. Ther. Med. 18. doi:10.3892/etm.2019.7619.

Malpeli, G., Innamorati, G., Decimo, I., Bencivenga, M., Kamdje, A. H. N., Perris, R., et al. (2019). Methylation dynamics of RASSF1A and its impact on cancer. Cancers (Basel). 11. doi:10.3390/cancers11070959.

Matsuoka, T., and Yashiro, M. (2016). Molecular targets for the treatment of pancreatic cancer: Clinical and experimental studies. World J. Gastroenterol. 22, 776–789. doi:10.3748/wjg.v22.i2.776.

Mazor, T., Pankov, A., Song, J. S., and Costello, J. F. (2016). Intratumoral Heterogeneity of the Epigenome. Cancer Cell 29, 440–451. doi:10.1016/j.ccell.2016.03.009.

McGranahan, N., and Swanton, C. (2017). Clonal Heterogeneity and Tumor Evolution: Past, Present, and the Future. Cell 168, 613–628. doi:10.1016/j.cell.2017.01.018.

Mishra, N. K., and Guda, C. (2017). Genome-wide DNA methylation analysis reveals molecular subtypes of pancreatic cancer. Oncotarget 8, 28990–29012. doi:10.18632/oncotarget.15993.

Mishra, N. K., Southekal, S., and Guda, C. (2019). Survival analysis of multi-omics data identifies potential prognostic markers of pancreatic ductal adenocarcinoma. Front. Genet. 10. doi:10.3389/fgene.2019.00624.

Moarii, M., Boeva, V., Vert, J. P., and Reyal, F. (2015). Changes in correlation between promoter methylation and gene expression in cancer. BMC Genomics 16. doi:10.1186/s12864-015-1994-2.

Munaron, L. (2015). Systems biology of ion channels and transporters in tumor angiogenesis: An omics view. Biochim. Biophys. Acta – Biomembr. 1848, 2647–2656. doi:10.1016/j.bbamem.2014.10.031.

Natale, F., Vivo, M., Falco, G., and Angrisano, T. (2019). Deciphering DNA methylation signatures of pancreatic cancer and pancreatitis. Clin. Epigenetics 11. doi:10.1186/s13148-019-0728-8.

Neureiter, D., Jäger, T., Ocker, M., and Kiesslich, T. (2014). Epigenetics and pancreatic cancer: Pathophysiology and novel treatment aspects. World J. Gastroenterol. 20, 7830–7848. doi:10.3748/wjg.v20.i24.7830.

Nones, K., Waddell, N., Song, S., Patch, A. M., Miller, D., Johns, A., et al. (2014). Genome-wide DNA methylation patterns in pancreatic ductal adenocarcinoma reveal epigenetic deregulation of SLIT-ROBO, ITGA2 and MET signaling. Int. J. Cancer 135, 1110–1118. doi:10.1002/ijc.28765.

Omura, N., and Goggins, M. (2009). Epigenetics and epigenetic alterations in pancreatic cancer. Int. J. Clin. Exp. Pathol. 2, 310–326. Available at: www.ijcep.com/IJCEP810007 [Accessed June 14, 2021].

Paradise, B. D., Barham, W., and Fernandez-Zapico, M. E. (2018). Targeting epigenetic aberrations in pancreatic cancer, a new path to improve patient outcomes? Cancers (Basel). 10. doi:10.3390/cancers10050128.

Patel, S. H., Edwards, M. J., and Ahmad, S. A. (2019). Intracellular ion channels in pancreas cancer. Cell. Physiol. Biochem. 53, 44–51. doi:10.33594/000000193.

Peng, Y., Wu, Q., Wang, L., Wang, H., and Yin, F. (2020). A DNA methylation signature to improve survival prediction of gastric cancer. Clin. Epigenetics 12. doi:10.1186/s13148-020-0807-x.

Ruscica, M., Dozio, E., Motta, M., and Magni, P. (2007). Relevance of the Neuropeptide Y System in the Biology of Cancer Progression. Curr. Top. Med. Chem. 7, 1682–1691. doi:10.2174/156802607782341019.

Saha, G., Singh, R., Mandal, A., Das, S., Chattopadhyay, E., Panja, P., et al. (2020). A novel hotspot and rare somatic mutation p.A138V, at TP53 is associated with poor survival of pancreatic ductal and periampullary adenocarcinoma patients. Mol. Med. 26. doi:10.1186/s10020-020-00183-1.

Storz, P., and Crawford, H. C. (2020). Carcinogenesis of Pancreatic Ductal Adenocarcinoma. Gastroenterology 158, 2072–2081. doi:10.1053/j.gastro.2020.02.059.

Tan, A. C., Jimeno, A., Lin, S. H., Wheelhouse, J., Chan, F., Solomon, A., et al. (2009). Characterizing DNA methylation patterns in pancreatic cancer genome. Mol. Oncol. 3, 425–438. doi:10.1016/j.molonc.2009.03.004.

Thompson, M. J., Rubbi, L., Dawson, D. W., Donahue, T. R., and Pellegrini, M. (2015). Pancreatic cancer patient survival correlates with DNA methylation of pancreas development genes. PLoS One 10. doi:10.1371/journal.pone.0128814.

Vedeld, H. M., Goel, A., and Lind, G. E. (2018). Epigenetic biomarkers in gastrointestinal cancers: The current state and clinical perspectives. Semin. Cancer Biol. 51, 36–49. doi:10.1016/j.semcancer.2017.12.004.

Vincent, A., Omura, N., Hong, S. M., Jaffe, A., Eshleman, J., and Goggins, M. (2011). Genome-wide analysis of promoter methylation associated with gene expression profile in pancreatic adenocarcinoma. Clin. Cancer Res. 17, 4341–4354. doi:10.1158/1078-0432.CCR-10-3431.

Wang, H., Guan, Q., Nan, Y., Ma, Q., and Zhong, Y. (2019). Overexpression of human MX2 gene suppresses cell proliferation, migration, and invasion via ERK/P38/NF-κB pathway in glioblastoma cells. J. Cell. Biochem. 120, 18762–18770. doi:10.1002/jcb.29189.

Xu, R., Xu, Q., Huang, G., Yin, X., Zhu, J., Peng, Y., et al. (2019). Combined Analysis of the Aberrant Epigenetic Alteration of Pancreatic Ductal Adenocarcinoma. Biomed Res. Int. 2019. doi:10.1155/2019/9379864.

Zhang, Q., Green, M. D., Lang, X., Lazarus, J., Parsels, J. D., Wei, S., et al. (2019). Inhibition of ATM increases interferon signaling and sensitizes pancreatic cancer to immune checkpoint blockade therapy. Cancer Res. 79, 3940–3951. doi:10.1158/0008-5472.CAN-19-0761.

Zouridis, H., Deng, N., Ivanova, T., Zhu, Y., Wong, B., Huang, D., et al. (2012). Methylation subtypes and large-scale epigenetic alterations in gastric cancer. Sci. Transl. Med. 4. doi:10.1126/scitranslmed.3004504.

