## Supplementary Material for "Altered DNA methylation in ion transport and immune signalling genes is associated with severity in Pancreatic Ductal Adenocarcinoma"

### 1 Supplemental Methods:

**Protocol for Expression measurement and analysis of GAPDH, ACTB, FAIM2, FOXE1, NPY, SLITRK3, SIGIRR, KCNA6 and RASSF1A genes in PDAC tumor tissues**

The expression of *GAPDH*, *ACTB,*  were analyzed in the Real Time PCR with an initial denaturation step of 95°C for 10 min (*CCND1*), 20 sec (*ACTB* and *GAPDH*) followed by 38 cycles of 95°C for 20 sec, and 60°C for 30 sec. A melting curve analysis step was carried out at the end of amplification step, consisting of denaturation at 95°C for 1 min, re-annealing at 55°C for 30 sec and a further denaturation at 95°C for 30 sec. The expression of *FAIM2* and *FOXE1,* were analyzed with annealing 57°C and 57.5°C for 30 sec respectively. The expression of *NPY, SLITRK3 KCNA6* and *RASSF1A,* were analyzed with annealing 58°C for 30 sec. The expression of *SIGIRR,* was analyzed with annealing 59°C for 30 sec. The other remaining steps for all seven genes were same as above. A melting curve analysis step was carried out at the end of amplification step, consisting of denaturation at 95°C for 1 min, re-annealing at 55°C for 30 sec and a further denaturation at 95°C for 30 sec. All primers for respective loci mentioned in supplementary Table 2

**Relative Gene Expression and Fold Change Analysis of Tumor Normal Paired Samples:**

The relative expression of seven genes namely *SLITRK3, FAIM2, FOXE1, NPY, RASSF1A, KCNA6,* and *SIGIRR* were studied in PDAC tumor with adjacent normal tissue samples. 2 genes, namely *GAPDH* and *ACTB* were used as internal control (or, reference genes) to normalize the expression of the target gene to compensate for any difference in the amount of sample tissue.

**Estimation of differential expression of genes in tumor and adjacent normal tissues:**

Target and reference gene Ct values were derived from the mean of duplicate. *GAPDH* and *ACTB* were used as reference control for normalization of target genes (Yu et al. 2015). Relative expression of targeted genes determined as 2^-ΔΔct^ was calculated for each of the samples to identify fold change. More than 2 fold change was identified as dysregulation (overexpressed/under expressed) for the respective genes.

**Statistical Analysis**

**Distribution of fold change differences and analysis of differential expression of genes in tumor and normal groups**

Distributions of 2^-Δct^ values after normalization with *GAPDH/ACTB* for the respective genes were checked by Anderson-Darling test in R for both matched tumor and adjacent normal groups (n=20) for the discovery group and (n=14) for the replicative group. Wilcoxon signed rank test was used to measure any significant differences (p≤0.05) of 2^-Δct^ values between two groups for all genes. Fold change differences between tumor and normal group of respective genes represented by box plots (ggplot2 package in R Studio).

### 2 Supplementary Figures

**Supplementary figure 1.**Schematic representation of the workflow : The step wise analysis pipeline with important cut-off factors and packages involved are mentioned in a systemic manner.

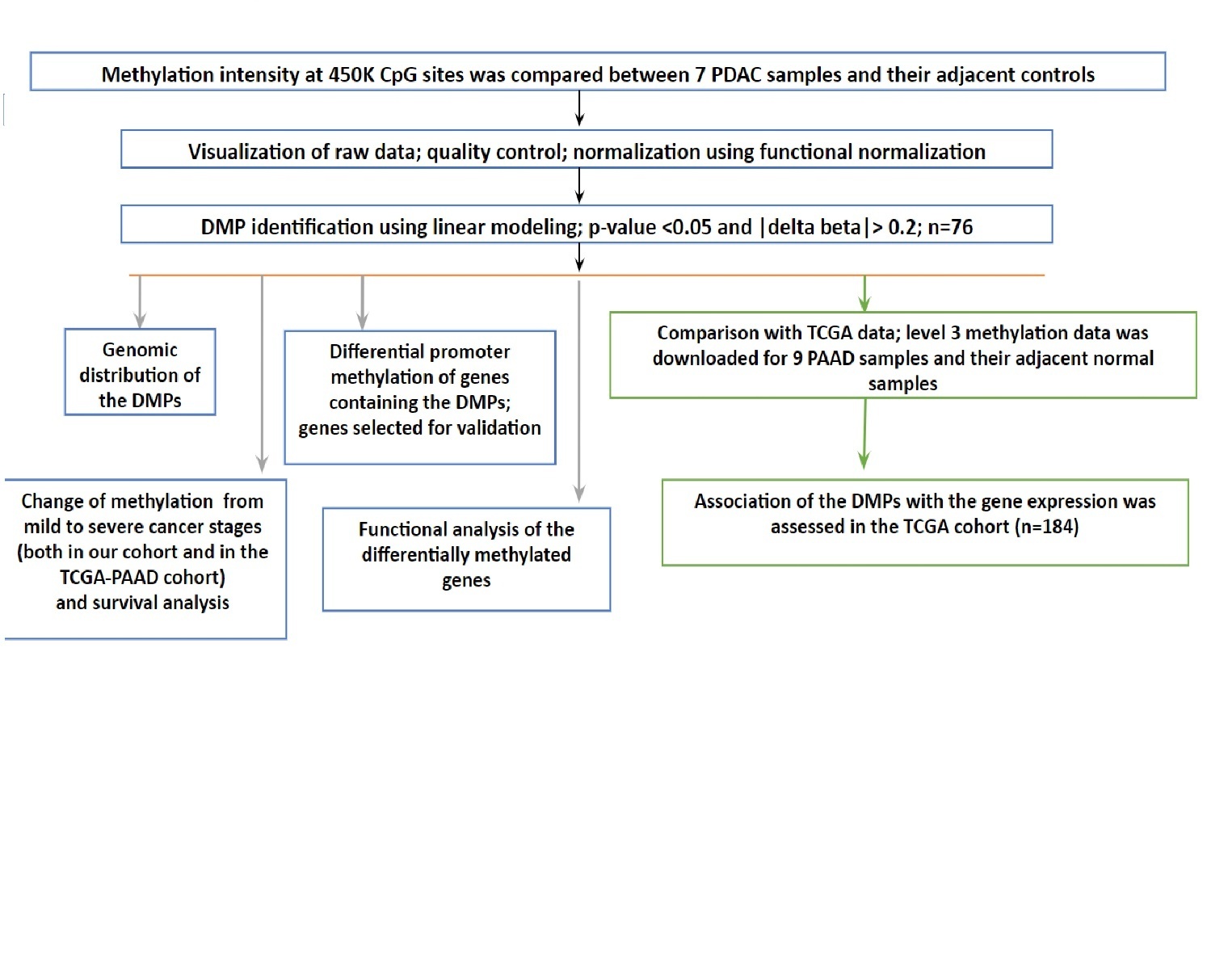

**Supplementary figure 2:** Target analysis of differential methylation at the 2213 DMPs (identified by Nones et al. ) in our study samples. Hierarchical clustering of the samples showed separate clustering of normal and cancer samples.

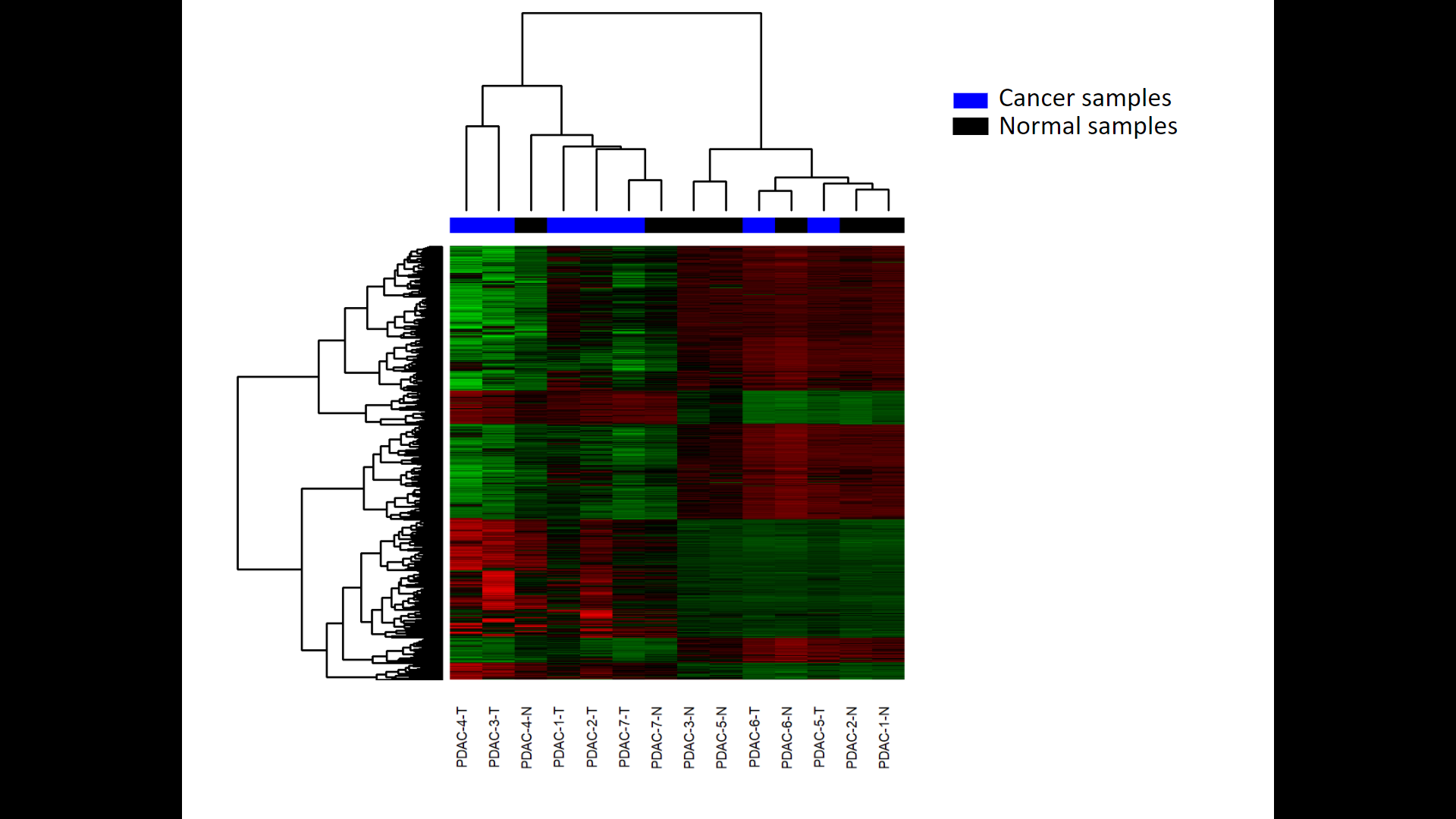

**Supplementary figure 3.** Boxplots showing distribution of average beta values at the differentially methylated positions (DMPs, n=76): (A) Hypomethylated DMPs (n=34).

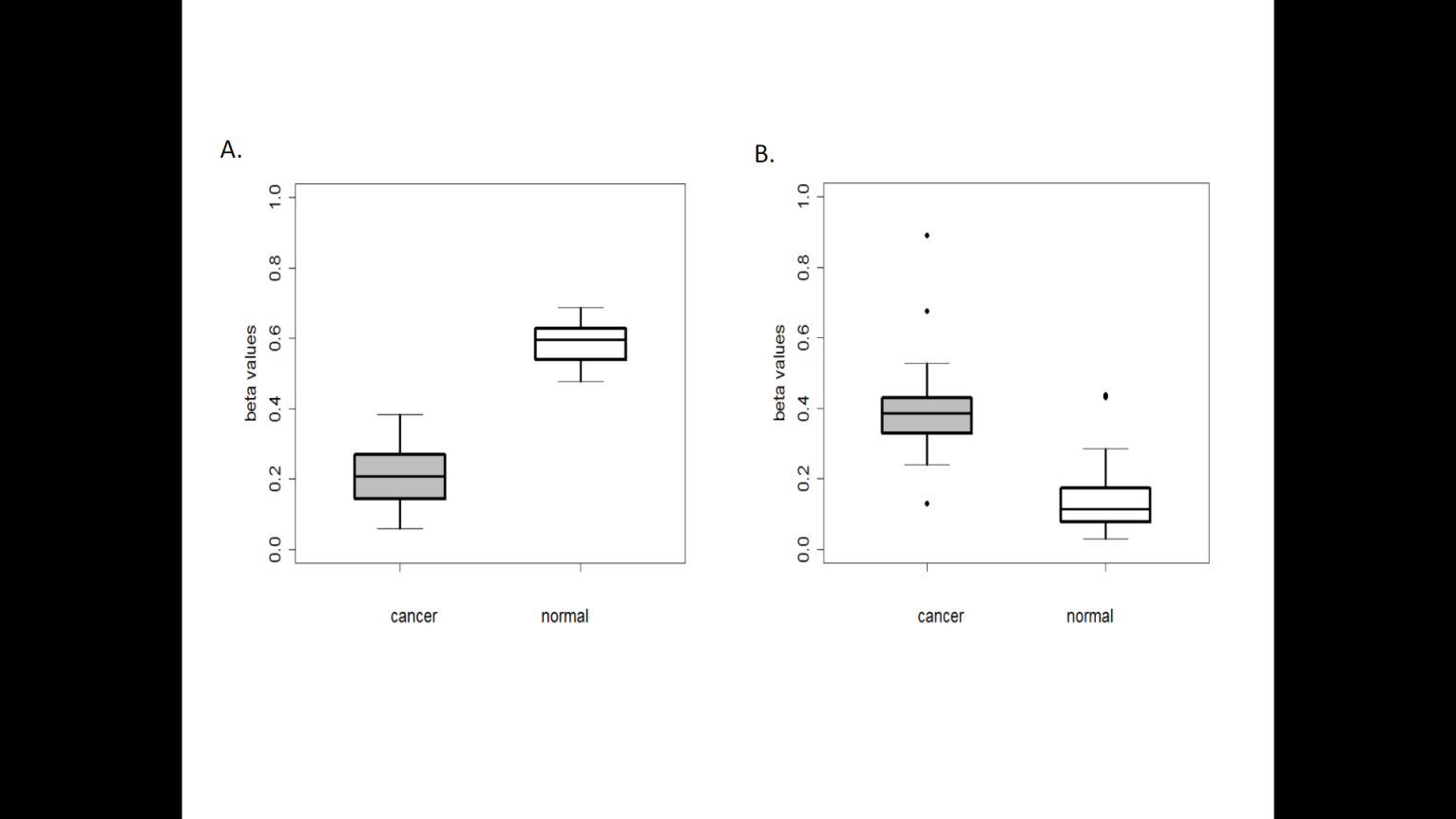

**Supplementary figure 4.** Validation of our study finding in the TCGA cohort. DNA methylation data from cancer and solid tissue normal samples was downloaded for 9 PAAD patients in the TCGA cohort. (A) Using our data analysis pipeline, 7832 DMPs were identified: 54% were hypermethylated and 46% were hypomethylated. (B) Hierarchical clustering analysis showed separate classification of cancer and solid tissue normal samples, based on 7832 DMPs. (C) Among the 7832 DMPs, 38 DMPs were common to our study finding. (D) Correlation plot between the beta values at these 38 DMPs in the TCGA data (for 9 PAAD patients) and in our study.

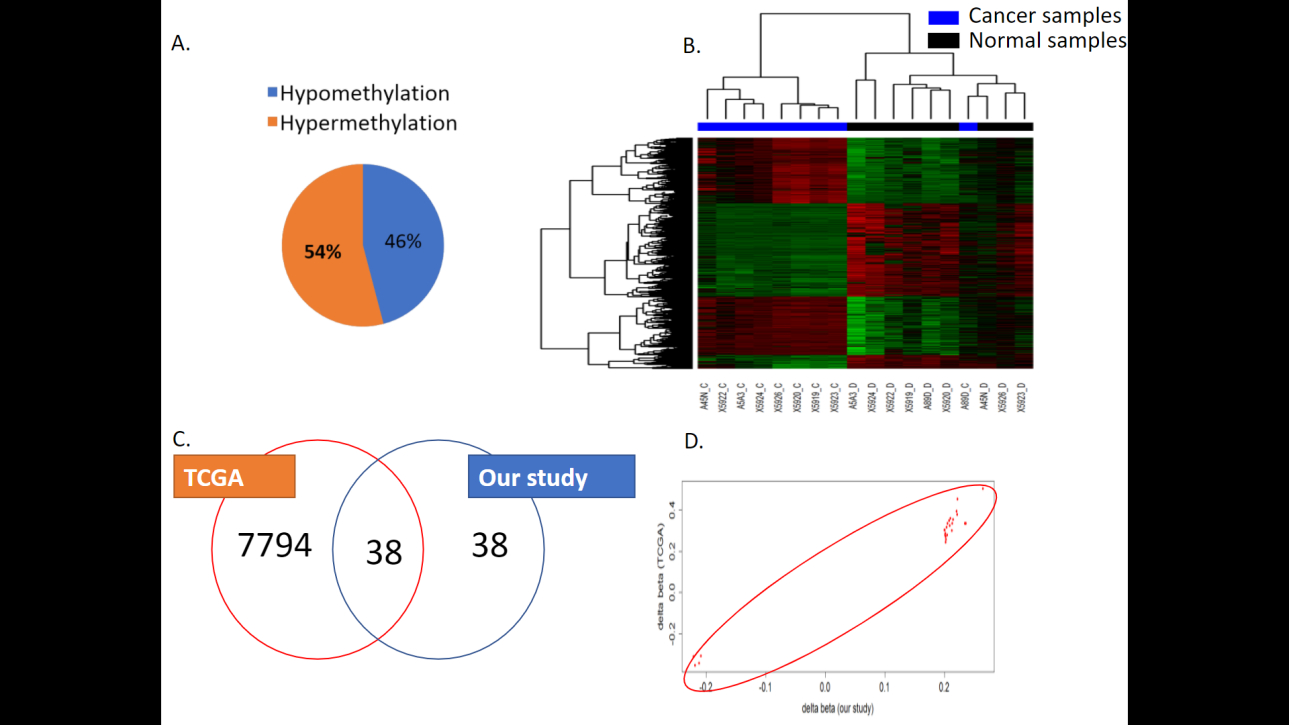

**Supplementary figure 5.** Distribution of average beta values at the promoters of 6 target genes between Tumor and Adjacent Normal.

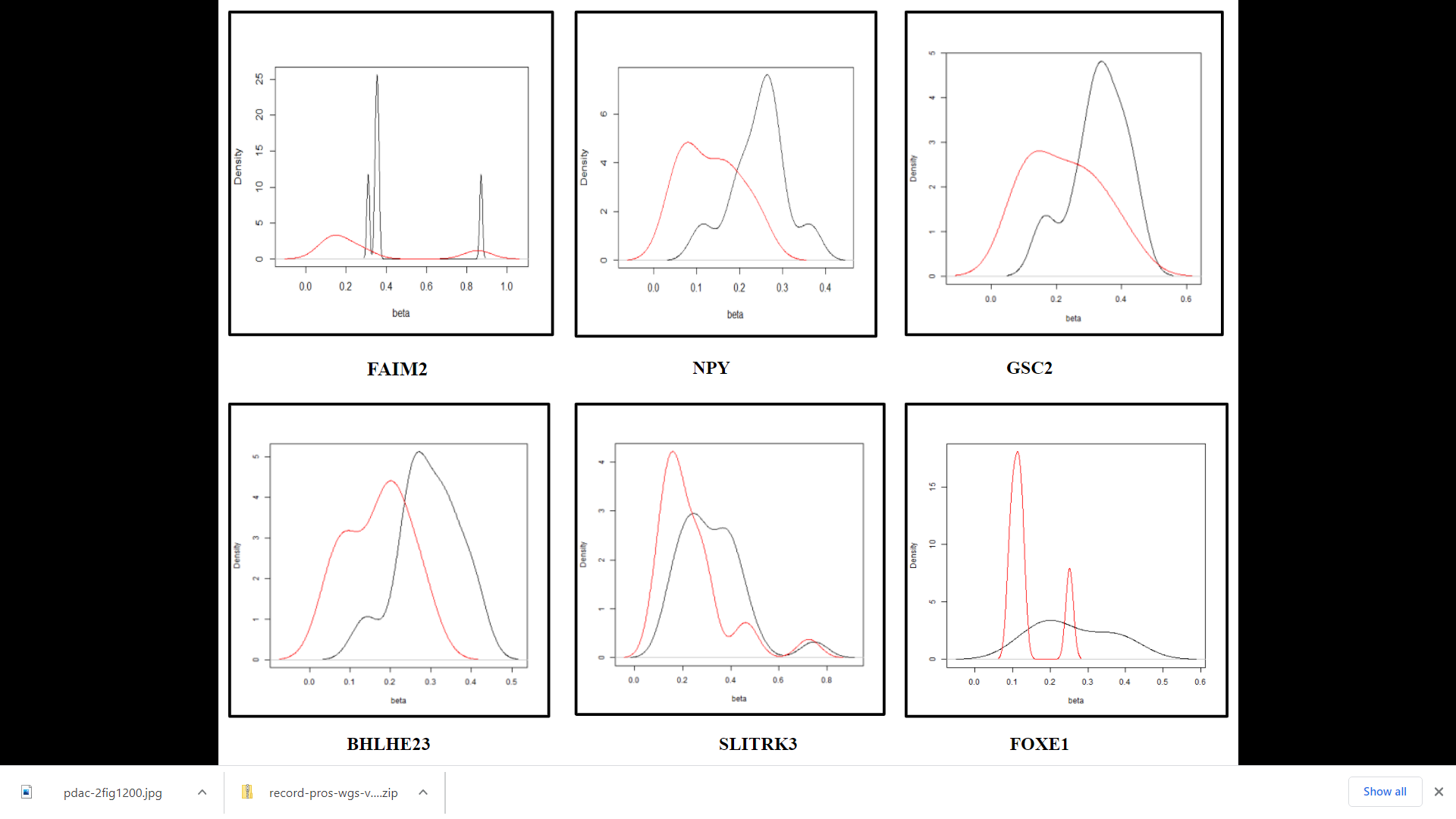

C3: Transcription regulators and other oncogenic proteins
